# Learning residue-level context for modeling protein-protein interactions

**DOI:** 10.64898/2026.06.01.729118

**Authors:** Zechuan Zhang, Zongxin Yang, Anbang Liu, Kun-Hsing Yu, Junhan Zhao, Yi Yang, Benjamin Neale, Siwei Chen

## Abstract

Protein language models (PLMs) enable prediction of protein properties by learning residue-level features from sequence, yet most PLM-based approaches to protein-protein interactions aggregate information across entire proteins, limiting resolution and interpretability. Here we present ReCLIP, a transformer-based framework that learns interaction-specific representations at the level of individual residues by combining intra-protein residue neighborhoods with residue-conditioned representations of interaction partners. We show that residue-centered context provides a general framework for modeling protein interactions across diverse biological settings. ReCLIP accurately predicts mutation-induced perturbations (AUROC = 0.973), generalizes to post-translational modifications that do not alter sequence (AUROC = 0.822), and enables zero-shot prediction of peptide-MHC binding across unseen alleles (AUROC up to 0.972). Analysis of learned residue neighborhoods reveals structurally and functionally coherent patterns aligned with known determinants of binding. Applied to clinically annotated genetic variants, ReCLIP identifies disease-associated interaction perturbations that link pathogenic variants to specific molecular interaction contexts. Our results establish a generalizable and interpretable framework for modeling protein interactions and provide insights into how residue-level context shapes interaction specificity and its perturbation.

## Introduction

Protein language models (PLMs) have achieved remarkable success in modeling protein sequence-function relationships. Trained on millions of natural proteins, these models learn representations (“embeddings”) that capture diverse biochemical and evolutionary properties directly from amino acid sequences, enabling prediction of structure, function, and variant effects for virtually any given protein.^1–3^

However, proteins rarely function in isolation, as most cellular processes are mediated by protein-protein interactions (PPIs). Modeling PPIs is inherently more complex than modeling individual proteins, because interaction outcomes depend not only on each interacting partner but also on context-dependent interactions between specific residues across proteins. Early PLM-based approaches to PPIs extended single-sequence models by concatenating partner sequences.^4–8^ Although concatenation captures compatibility between partners, it treats an interacting protein pair as a single unified entity, potentially obscuring interaction-specific features and degrading representation quality. More recent methods introduced interaction-aware architectures that learn partner-conditioned representations of proteins, leading to improved performance in PPI-related tasks.^9,10^ Nevertheless, most existing approaches still summarize each protein or protein pair into a fixed embedding, creating a resolution bottleneck in which residue-specific determinants of binding can be diluted before prediction.

A key biological insight is that PPIs are often governed by a restricted subset of residues.^11,12^ Large-scale structural and genetic studies have shown that mutations perturbing PPIs disproportionately localize to interaction interfaces, modulating local binding without globally altering protein structure or stability.^13–16^ These observations underscore that PPI function is encoded within localized residue neighborhoods rather than distributed uniformly across entire sequences. This raises a fundamental question: which residue neighborhoods are most informative for modeling interaction outcomes?

Sequence adjacency provides a simple definition of neighborhood and has been adopted in previous interaction modeling approaches.^17^ However, residues distant in sequence can become proximal in three-dimensional space upon protein folding, forming functional neighborhoods that are not captured by sequence-based adjacency.^18–20^ Advances in structure prediction, notably through AlphaFold,^21^ have enabled inference of structural neighborhoods directly from sequence. Yet accurate prediction of multi-protein structures remains infeasible at the scale of hundreds of thousands of PPIs, and experimentally resolved complex structures are available for only a small fraction of known interactions.^22^ Even when structures are available, spatial proximity alone does not fully capture functional dependencies driven by allosteric coupling, conformational dynamics, and other long-range effects.^22,23^ These limitations motivate a learned definition of residue context that is not restricted to sequence adjacency or predefined structural distance.

In this study, we present Residue-level Context Learning for Interacting Proteins (ReCLIP), a transformer-based interaction modeling framework that learns interaction-specific, residue-level dependencies from PLM representations. ReCLIP models PPIs through learning functional residue neighborhoods from internal PLM attention patterns, without restricting context to sequence proximity or explicit structural information. By organizing interaction modeling around a specific residue, ReCLIP constructs residue-centered representations that capture how local context within one protein and residue-conditioned dependencies with its partner jointly determine interaction outcomes. This formulation reframes PPI prediction as a residue-centered interaction modeling problem rather than a protein-level embedding classification task.

We validated ReCLIP in diverse biological settings, where it consistently outperforms state-of-the-art methods. First, ReCLIP accurately predicts mutation-induced perturbations to PPIs across multiple interaction outcome classes (area under the receiver operating characteristic curve [AUROC] = 0.973, *P*<1e-200). Second, ReCLIP generalizes to post-translational modifications (PTMs) that regulate interactions without changing the primary sequence, a setting not accessible to methods that require explicit wild-type versus mutant comparisons (AUROC = 0.822, *P* = 3.48e-39). Third, ReCLIP effectively models peptide-MHC (pMHC) interactions, a clinically important class of PPIs characterized by extensive allelic diversity (AUROC up to 0.972; *P* = 1.02e-9). Analysis of the learned residue neighborhoods reveals structurally and functionally coherent contexts associated with improved predictive performance. Finally, systematic application to over 140,000 clinically annotated variants (ClinVar^24^) identifies interaction perturbations enriched among pathogenic variants and organized within disease-relevant molecular contexts. Together, these results establish a generalizable and interpretable framework for modeling PPIs and for understanding how residue-level context encodes interaction specificity, its perturbation, and the molecular consequences of human genetic variation.

## Results

### Overview of the ReCLIP framework

PLMs learn context-dependent representations of individual residues that capture biochemical, structural, and evolutionary constraints encoded in protein sequences.^1–3^ Yet the use of such residue-level representations for modeling PPIs remains limited by how they are summarized for prediction. Most PLM-based approaches to PPIs, including sequence-concatenation or interaction-aware architectures, ultimately represent each protein or protein pair with a fixed embedding (**Fig. 1a**).^4–10^ This protein-level aggregation is misaligned with the biology of protein interactions, which are often governed by a restricted subset of residues that mediate binding and determine specificity.^11,12^ As a result, residue-level determinants of interaction outcomes can be diluted when information is pooled across entire sequences.

**Figure 1.**
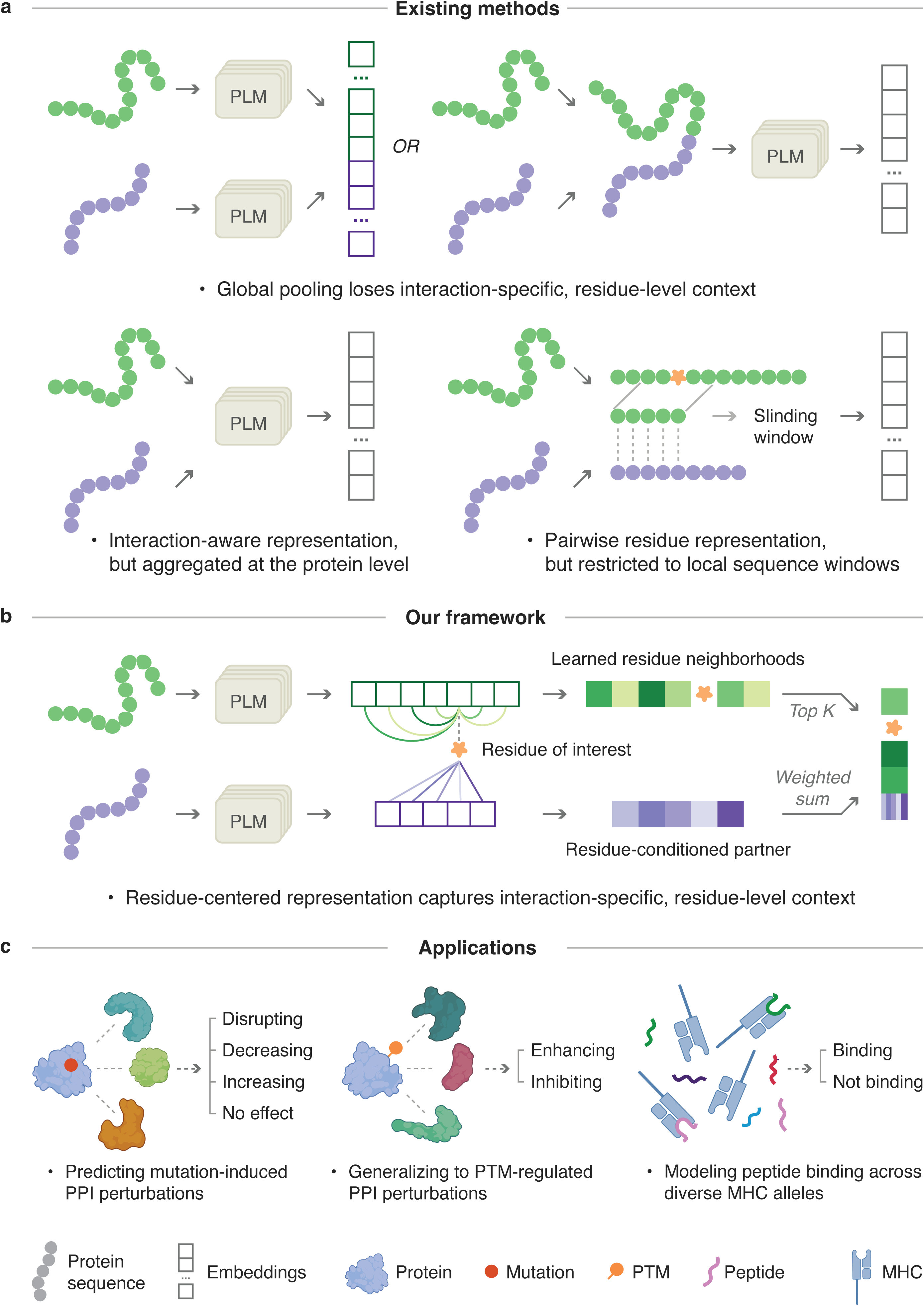
Overview of ReCLIP for residue-centered modeling of protein-protein interactions (PPIs). **a**, Limitations of existing protein language model (PLM)-based approaches for PPI modeling. Current methods often rely on protein-level aggregation or local sequence windows, which can obscure residue-level interaction context or miss long-range dependencies. **b**, Overview of the ReCLIP framework. ReCLIP organizes interaction representations around residue-centered context, combining learned intra-protein functional neighborhoods with residue-conditioned representations of interaction partners to capture interaction-specific, residue-level information. **c**, Applications of ReCLIP across diverse PPI prediction tasks. Left, prediction of mutation-induced perturbations with four outcome classes (disrupting, decreasing, increasing, no effect). Middle, prediction of PTM-regulated interaction changes (enhancing or inhibiting). Right, prediction of peptide-MHC (pMHC) binding across diverse MHC alleles.

To address this gap, we developed Residue-level Context Learning for Interacting Proteins (ReCLIP), a framework that formulates PPI modeling as a residue-centered, context-dependent problem rather than a protein-level classification task. Conceptually, ReCLIP asks: given a specific residue on a protein, which residues within that protein and on its binding partner are most informative for the interaction outcome? By explicitly centering representation learning on a queried residue, ReCLIP constructs a residue-specific interaction context rather than assigning a single fixed representation to the entire protein pair.

ReCLIP employs the attention mechanism of PLMs to construct query-centered context (Methods). Given an interacting protein pair (A-B) and a residue of interest on protein A, ReCLIP first defines a functional neighborhood as a small set of residues in protein A prioritized by attention from the queried residue (**Fig. 1b**). This attention-defined neighborhood allows ReCLIP to capture residue dependencies that may be nearby in sequence, proximal in three-dimensional structure, or functionally coupled through longer-range constraints.

Second, ReCLIP builds a residue-conditioned representation of the interaction partner. Because not all residues on the partner (i.e., protein B) contribute to binding equally, ReCLIP computes cross-protein attention from the queried residue to partner residues and uses these weights to define an interaction relevance distribution (Methods). The partner residue embeddings are then aggregated according to this distribution, producing a representation of protein B that is explicitly conditioned on the queried residue rather than on the interaction pair as a whole (**Fig. 1b**). This design enables the same protein pair to be represented differently depending on the queried residue, supporting a many-to-many mapping between query and partner residues without requiring explicit structural annotations or predefined interfaces.

Finally, ReCLIP combines the residue-centered intra-protein context and the residue-conditioned partner representation to form a joint interaction representation (**Fig. 1b**). This representation is used for predicting interaction specificity and quantitative changes in binding driven by residue-level perturbations (**Fig. 1c**). By constructing representations around residue-specific context prior to prediction, ReCLIP captures interaction-specific, residue-level information that would otherwise be lost through early global pooling.

### ReCLIP predicts mutation-induced perturbations to PPIs

We first evaluated ReCLIP on predicting mutation-induced perturbations to PPIs, a biologically important task as many functional missense mutations act by perturbing specific protein interactions rather than overall protein stability.^13–16^ We used the IntAct database,^25^ which curates experimentally characterized missense mutations with annotated effects on protein binding, comprising 19,559 mutations evaluated across 56,429 PPIs (Methods). We formulated this as a four-class classification problem, distinguishing mutations that disrupt, decrease, increase, or do not affect binding strength (**Fig. 2a**). For each mutation, the model takes as input the sequence of the mutated protein, the mutation position, and the sequence of its interaction partner. Predictive performance was evaluated on held-out test data under stratified cross-validation (Methods).

**Figure 2.**
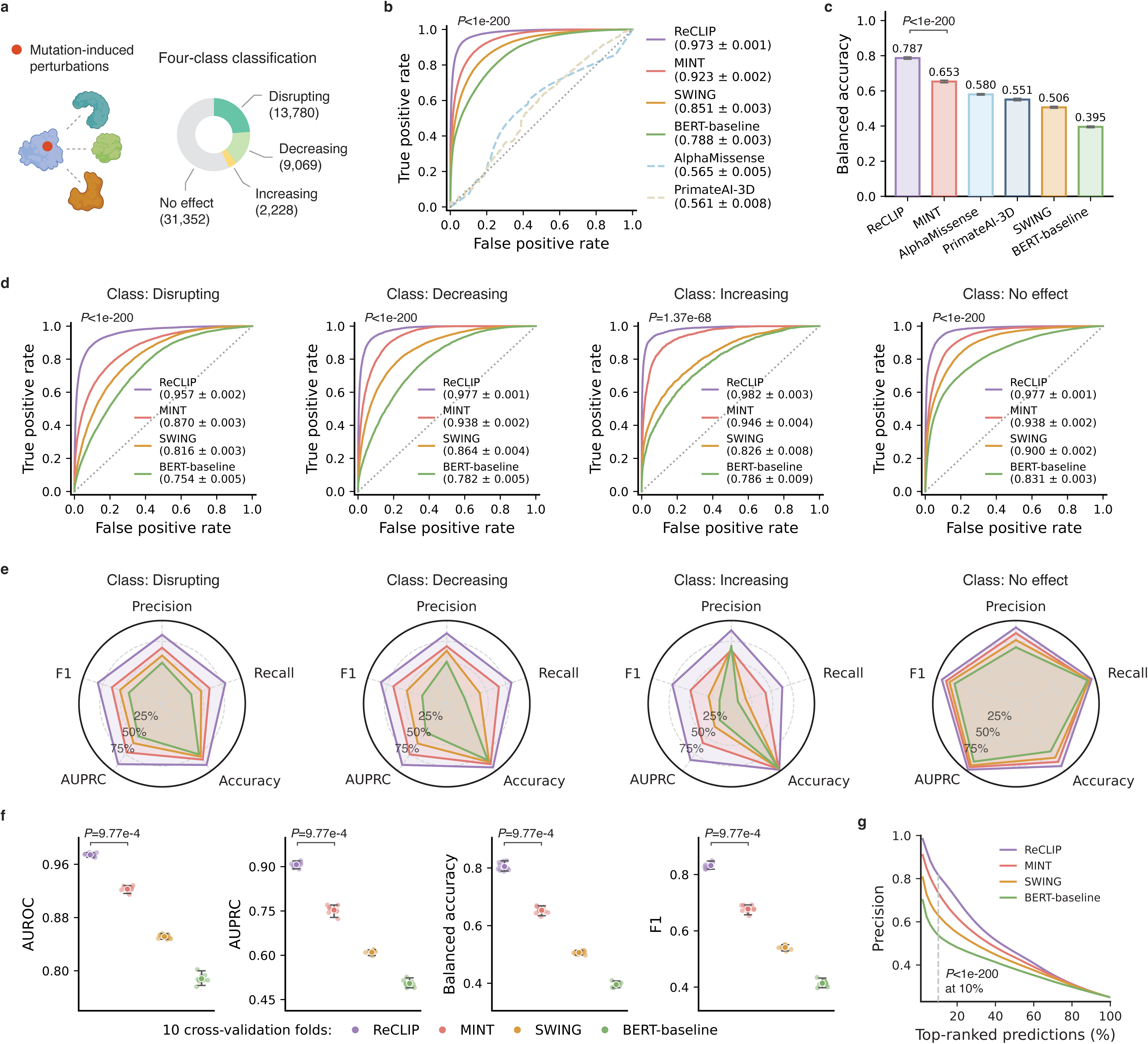
ReCLIP predicts mutation-induced perturbations to PPIs. **a**, Overview of the mutation-induced interaction perturbation prediction, formulated as a four-class classification problem (disrupting, decreasing, increasing, no effect). **b**, Receiver operating characteristic (ROC) curves comparing ReCLIP with baseline methods across all mutation classes. ReCLIP achieves the highest overall AUROC. **c**, Balanced accuracy comparison across methods, highlighting ReCLIP’s robustness to class imbalance. **d**, Class-specific ROC curves for each interaction outcome. ReCLIP consistently outperforms comparison methods across all classes. **e**,**f**, Performance comparison across evaluation metrics for each prediction class (**e**) and across 10 cross-validation folds (**f**). ReCLIP shows high and consistent performance across these settings. In **f**, each point represents one fold; center lines indicate medians and error bars denote the 2.5^th^-97.5^th^ percentile range. Statistical significance was assessed using a two-sided Wilcoxon signed-rank test. **g**, Precision as a function of top-ranked predictions. ReCLIP maintains higher precision across the ranking range, exceeding 80% at the top 10% threshold. Statistical significance was assessed using bootstrap resampling (n = 200) of the difference in precision between ReCLIP and the second-best method. For **b-d**, evaluation metrics are reported as mean with 95% confidence intervals (CIs) computed from 200 bootstrap resamples; statistical significance was assessed from the bootstrap distribution of metric differences between ReCLIP and the second-best method.

ReCLIP achieved an overall AUROC of 0.973, outperforming several representative classes of recent approaches (bootstrap *P*<1e-200 versus second-best; **Fig. 2b**). These include an interaction-aware model that encodes protein pairs into a single embedding (MINT,^9^ AUROC = 0.923), a sequence-adjacency-based approach that restricts residue context to local windows (SWING,^17^ AUROC = 0.851), and a global embedding baseline that operates on pooled protein representations (BERT-baseline, AUROC = 0.788). PPI-unaware variant effect predictors, such as AlphaMissense^26^ and PrimateAI-3D,^27^ showed substantially lower performance in this setting (AUROCs = 0.564 and 0.561, respectively), consistent with their focus on single-protein-level variant effects rather than interaction-specific outcomes.

To account for the class imbalance arising from the overrepresentation of no-effect variants (56.8%; **Fig. 2a**), we further evaluated performance using balanced accuracy. ReCLIP achieved a balanced accuracy of 0.787, exceeding the second-best model (MINT, 0.653) by 0.134 (bootstrap *P*<1e-200; **Fig. 2c**). Performance of sequence-adjacency-based and global embedding approaches decreased more markedly (0.506 and 0.395, respectively), suggesting reduced robustness to class imbalance. Moreover, ReCLIP maintained strong discrimination across individual interaction outcome classes (AUROCs of disrupting = 0.957 [*P*<1e-200], decreasing = 0.977 [*P*<1e-200], increasing = 0.982 [*P* = 1.37e-68], and no-effect mutations = 0.977 [*P*<1e-200]; **Fig. 2d**). The ranking of comparison methods remained consistent across classes, with the largest performance differences observed for the disrupting class. Across additional evaluation metrics, including accuracy, precision, recall, F1-score, and precision-recall AUC (AUPRC), ReCLIP consistently outperformed other models (**Fig. 2e**), and improvements were stable across cross-validation folds (Wilcoxon signed-rank test *P* = 9.77e-4; **Fig. 2f**).

Many existing mutation-effect prediction approaches focus on interaction disruption effects, we reformulated the task using only disrupting and no-effect labels and compared ReCLIP with additional PLM-based interaction models designed for binary classification.^28,29^ ReCLIP continued to outperform all comparison methods under this setting (**Extended Data Fig. 1**), demonstrating that its advantage generalizes across alternative formulations of interaction perturbation prediction.

Because functional validation is often limited to a small number of candidate mutations, we next evaluated ranking performance by ordering predictions according to model confidence and calculating macro precision across Top-K thresholds. ReCLIP maintained higher precision across the full ranking range compared to other methods (**Fig. 2g**). At the top 10% of predictions, precision reached 81.8% (bootstrap *P*<1e-200 versus second-best), indicating strong enrichment of correctly classified mutations among top-ranked candidates and supporting their prioritization for downstream functional studies.

### ReCLIP generalizes to PTM-regulated interaction perturbations

We then asked whether ReCLIP generalizes beyond sequence variation to regulatory perturbations, focusing on post-translational modifications (PTMs). This setting differs fundamentally from mutation-based prediction because PTMs do not alter the underlying protein sequence. Methods that rely on comparisons between wild-type and mutant sequences (e.g., MINT, AlphaMissense, and PrimateAI-3D) are therefore not applicable in this context.

We used the PTMint database,^30^ a manually curated collection of 3,052 experimentally supported PTM-regulated PPIs annotated as either enhancing or inhibiting binding (**Fig. 3a**; Methods). We formulated this as a binary classification problem and evaluated performance using the same cross-validation scheme as in the mutation-based prediction setting. Each instance is defined by the sequences of the interacting proteins and the position of the modification site (Methods).

**Figure 3.**
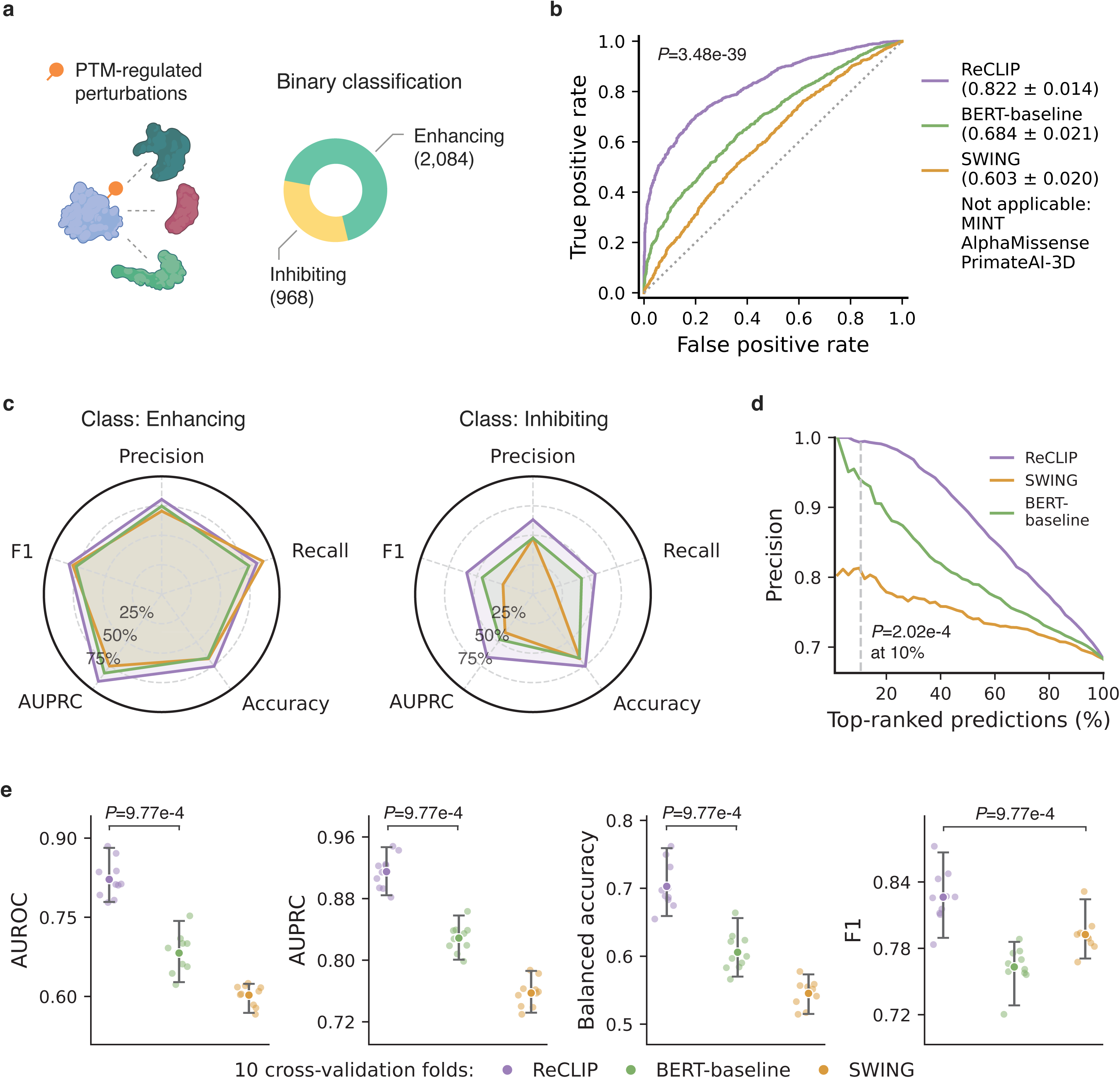
ReCLIP generalizes to PTM-regulated interaction perturbations. **a**, Overview of the PTM-regulated interaction perturbation prediction, formulated as a binary classification problem (enhancing versus inhibiting). **b**, ROC curves comparing ReCLIP with baseline methods. ReCLIP achieves the highest overall AUROC. AUROC values are reported as mean with 95% CIs computed from 200 bootstrap resamples; statistical significance was assessed from the bootstrap distribution of metric differences between ReCLIP and the second-best method. **c**,**d**, Performance comparison across evaluation metrics for each interaction outcome class (**c**) and across cross-validation folds (**d**). ReCLIP shows consistently strong performance across metrics and folds relative to baselines. In **d**, each point represents one fold; center lines indicate medians and error bars denote the 2.5^th^-97.5^th^ percentile range. Statistical significance was assessed using a two-sided Wilcoxon signed-rank test. **e**, Precision as a function of top-ranked predictions. ReCLIP maintains higher precision across the ranking range, remaining above 90% through the top 50% of predictions. Statistical significance was assessed using bootstrap resampling (n = 200) of the difference in precision between ReCLIP and the second-best method.

ReCLIP achieved an AUROC of 0.822, substantially outperforming comparison methods (bootstrap *P* = 3.48e-39 versus second-best; **Fig. 3b**). Improvements were consistent across evaluation metrics, cross-validation folds, and interaction outcome classes (**Fig. 3c,d**). Notably, the performance gain was larger for the underrepresented “inhibiting” class, highlighting ReCLIP’s robustness to class imbalance and limited sample sizes. ReCLIP also maintained high precision among top-ranked predictions, exceeding 90% through the top 50% threshold, whereas other methods showed a more pronounced decline (bootstrap *P* = 2.02e-4 at top 10%; **Fig. 3e**). Across evaluation metrics, the relative improvement over competing models was greater than that observed in the mutation-prediction setting. This pattern suggests that residue-centered modeling is especially effective when interaction outcomes must be inferred from subtle regulatory perturbations without explicit sequence changes.

### ReCLIP models pMHC binding across diverse MHC alleles

We further applied ReCLIP to predict pMHC binding, a distinct class of interactions involving short peptide ligands and highly polymorphic receptors.^31^ This setting is particularly challenging, as it requires generalization across diverse MHC alleles, including those not observed during training.

We therefore adopted a zero-shot evaluation framework in which test alleles were strictly excluded from the training set (**Fig. 4a**). We used curated datasets previously assembled for pMHC modeling,^17,32^ comprising 171,560 pMHC interaction pairs spanning 14 MHC alleles with varying degrees of functional similarity (**Fig. 4a,b**). For each pMHC pair, the peptide sequence and its central position were treated as the query and the MHC binding-groove region as the interaction partner.

**Figure 4.**
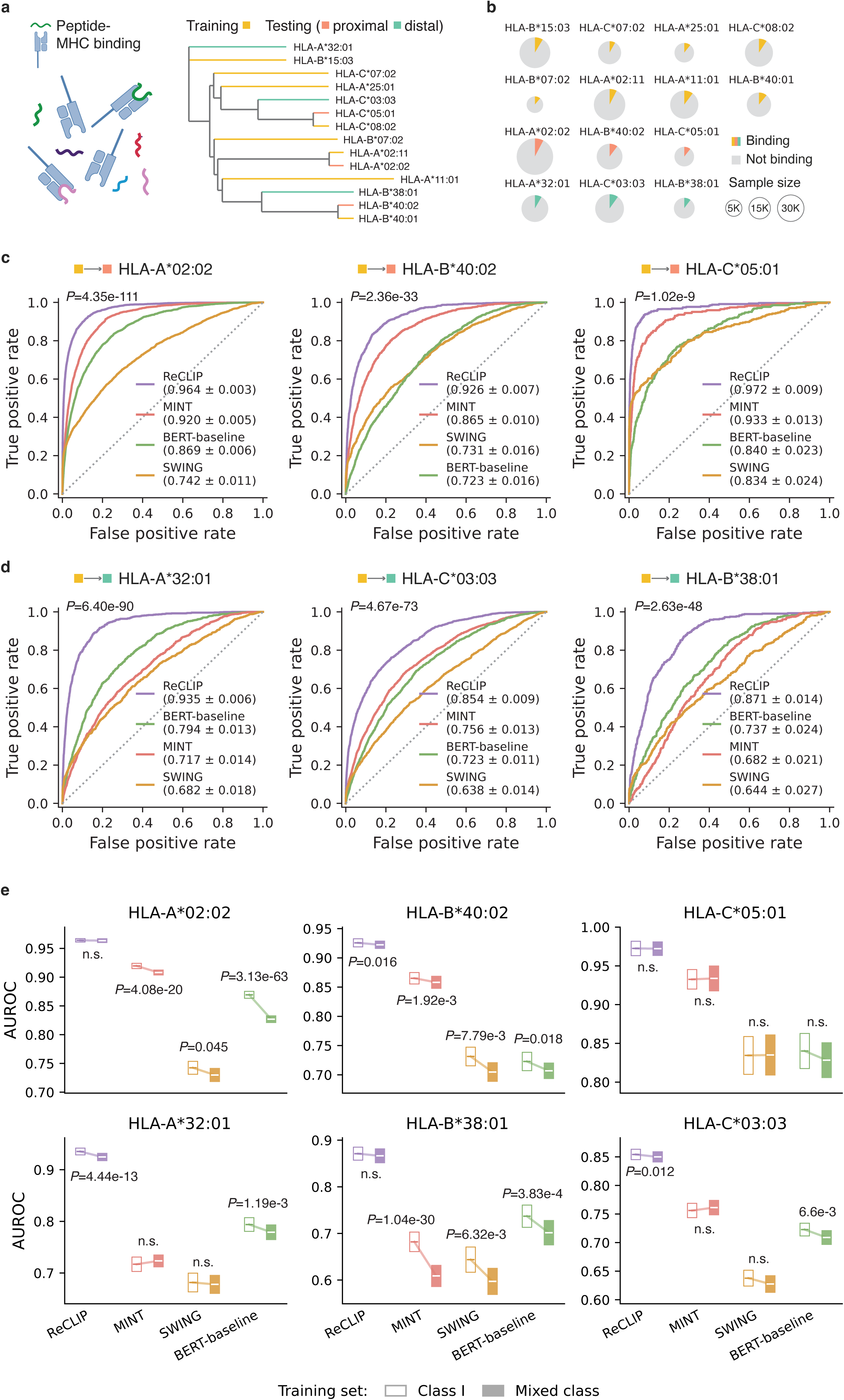
ReCLIP models peptide-MHC (pMHC) binding across diverse MHC alleles. **a**, Overview of the pMHC binding prediction task and zero-shot evaluation framework. MHC alleles are partitioned into training and held-out test sets such that test alleles are excluded from training. A hierarchical clustering tree based on functional similarity is used to define proximal and distal test alleles, enabling evaluation under varying degrees of distribution shift. **b**, Data composition across MHC alleles. Each circle represents one allele; circle size indicates sample size and color denotes binding (colored) versus non-binding (grey) composition. Sample sizes and binding class fractions are as follows: , HLA-B*15:03 (n=20,164; 8.6%), HLA-C*07:02 (n=6,855; 8.1%), HLA-A*25:01 (n=3,124; 10.7%), HLA-C*08:02 (n=14,574; 9.2%), HLA-B*07:02 (n=1,114; 11.1%), HLA-A*02:11 (n=21,499; 7.5%), HLA-A*11:01 (n=14,858; 10.3%), HLA-B*40:01 (n=8,486; 10.9%), HLA-A*32:01 (n=12,427; 8.3%), HLA-C*03:03 (n=15,954; 9.9%), HLA-B*38:01 (n=4,248; 10.3%), HLA-C*05:01 (n=3,193; 10.7%), HLA-A*02:02 (n=34,347; 7.7%), and HLA-B*40:02 (n=10,717; 10.3%). **c**,**d**, ROC curves for zero-shot prediction on functionally proximal (**c**) and distal (**d**) MHC alleles. ReCLIP achieves the highest performance across all held-out alleles. **e**, Performance comparison under class I (hollow boxes) and mixed-class (filled boxes) training. ReCLIP maintains the highest performance with greater stability across training settings. For **c-e**, AUROC values are reported as mean with 95% CIs computed from 200 bootstrap resamples. Statistical significance was assessed using the bootstrap distribution of metric differences between ReCLIP and the second-best method (**c**,**d**) or between training settings (**e**). n.s., not significant (*P* > 0.05).

We first trained the model on MHC class I alleles and evaluated it on three unseen alleles that were functionally proximal to the training set (HLA-A*02:02, HLA-B*40:02, and HLA-C*05:01). ReCLIP achieved AUROCs of 0.964, 0.926, and 0.972, respectively, outperforming all comparison methods across each held-out allele (bootstrap *P* = 4.35e-111, 2.36e-33, and 1.02e-9, respectively; **Fig. 4c**). To assess performance under stronger distribution shift, we evaluated three functionally distal alleles (HLA-A*32:01, HLA-C*03:03, and HLA-B*38:01). ReCLIP remained the top-performing model, achieving AUROCs of 0.935, 0.854, and 0.871, respectively, whereas competing methods declined more sharply, with average AUROCs falling below 0.75 (**Fig. 4d**).

We further examined model robustness by training a mixed-class model incorporating both class I and class II pMHC interactions and evaluating performance on the same six unseen class I alleles. ReCLIP maintained the highest performance with minimal performance degradation (ΔAUROC < 0.01; **Fig. 4e**), demonstrating robustness to increased interaction diversity. Together, these results suggest that residue-centered interaction modeling generalizes across diverse interaction contexts while preserving sensitivity to residue-level determinants of binding specificity.

### ReCLIP captures biologically meaningful residue neighborhoods

A key technical design of ReCLIP is that it represents PPIs through learned residue-centered context rather than pooled global embeddings or fixed sequence windows. To characterize the biological properties of these learned representations, we analyzed the mutation dataset at the level of individual residues. For each queried residue (mutation site), we defined its functional neighborhood as the five residues receiving the highest attention weights in each of the 20 attention heads and compared them with matched sets of randomly sampled residues from the same protein (Methods).

Functional neighborhoods were strongly localized in both sequence and structure. Relative to random controls, these residues were significantly closer to the queried residue along the primary sequence (medians = 16.7 versus 132.5 amino acids, Wilcoxon signed-rank test *P*<1e-200; **Fig. 5a**) and in three-dimensional space (medians = 11.5 versus 34.6 Å, Wilcoxon signed-rank test *P*<1e-200; **Fig. 5b**; Methods). Notably, this structural proximity remained evident even when restricting the analysis to residues distal in primary sequence (separated by more than 10 amino acids; medians = 19.6 versus 35.9 Å, Wilcoxon signed-rank test *P*<1e-200; **Fig. 5b**), indicating that the model captures non-local contacts arising from protein folding rather than simply recovering adjacent sequence contexts.

**Figure 5.**
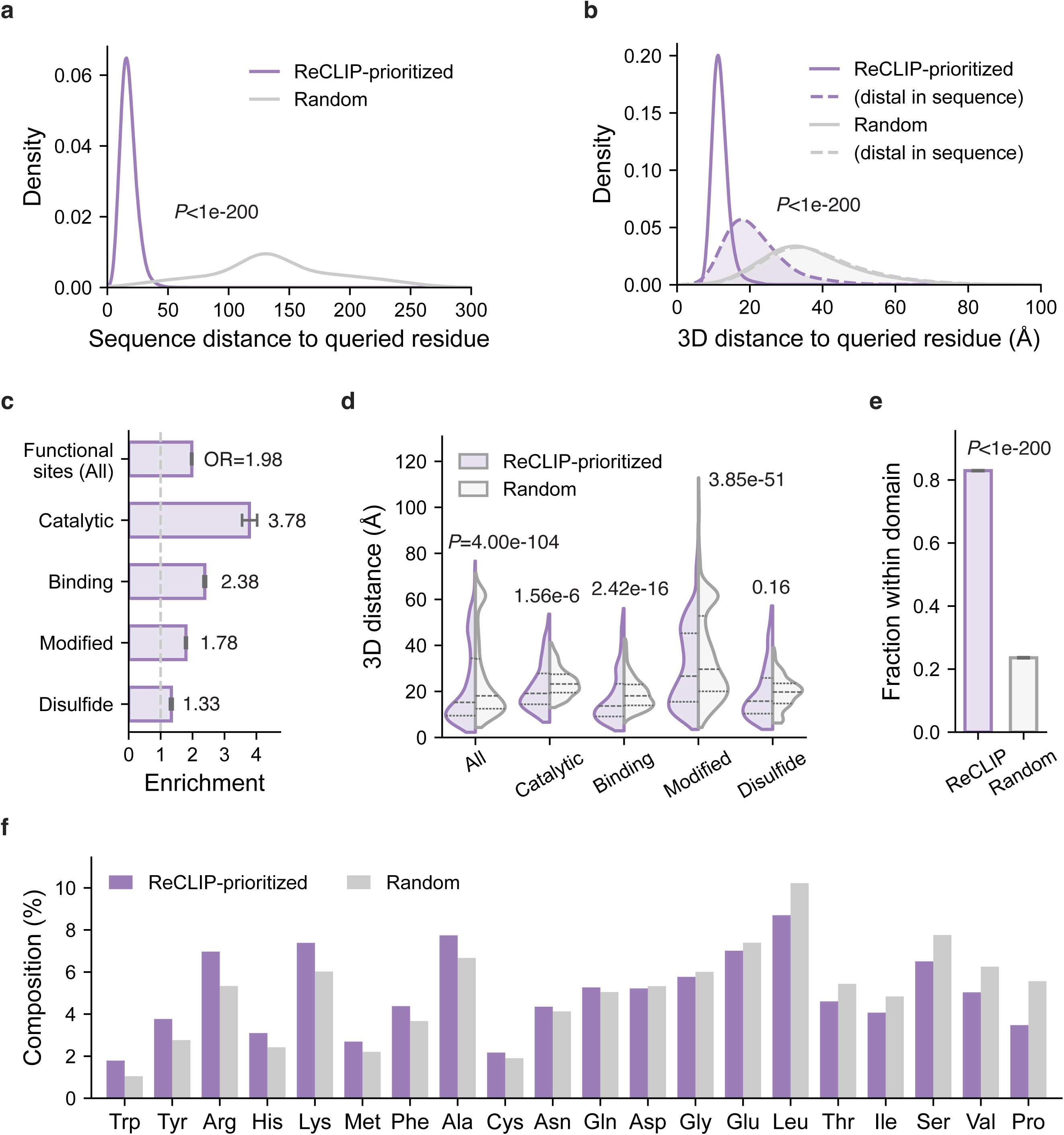
ReCLIP captures biologically meaningful residue neighborhoods. **a,b**, Distribution of sequence (**a**) and three-dimensional (**b**) distance between residues prioritized by ReCLIP (purple) and the queried residue, compared with randomly sampled residues from the same protein (grey). In **b**, dashed lines indicate residues that are distal in sequence (>10 amino acids), highlighting enrichment of non-local structural neighborhoods. **c**, Enrichment of prioritized residues in annotated functional sites, including catalytic residues, binding sites, modified residues, and disulfide bonds. Enrichment is quantified as odds ratios (ORs) relative to random expectation; error bars indicate 95% CIs. The dashed vertical line indicates an OR of 1 (no enrichment). **d**, Distribution of three-dimensional distances between prioritized residues (purple) and annotated functional sites, compared with random controls (grey). Dashed horizontal lines indicate medians and interquartile ranges. **e**, Fraction of prioritized residues (purple) located within the same annotated protein domain as the queried residue, compared with randomly sampled residues (grey). Error bars indicate Wilson 95% binomial CIs. **f**, Amino-acid composition of prioritized residues (purple) compared with random background (grey). Bars are ordered by relative enrichment compared to background. For **a-f**, statistical significance was assessed using two-sided Wilcoxon signed-rank tests (**a,b,d**) or Fisher’s exact tests (**c,e,f**).

We next evaluated whether these neighborhoods correspond to biologically meaningful regions. Residues within functional neighborhoods were significantly enriched for annotated functional sites,^33^ including catalytic residues, binding sites, modified residues, and disulfide bonds (odds ratios (ORs) = 1.33-3.78, all Fisher’s exact test *P*<1e-50; **Fig. 5c**; Methods). Consistently, these residues were located closer to functional sites in three-dimensional protein structures, with the strongest enrichment observed for binding sites (medians = 14.3 versus 19.2 Å, Wilcoxon signed-rank test *P*=2.42e-16; **Fig. 5d**). These neighborhoods also exhibited domain-level coherence, with a higher fraction of residues falling within the same annotated domain^34^ as the queried residue than expected by chance (0.83 versus 0.24 , Fisher’s exact test *P*<1e-200; **Fig. 5e**; Methods).

Moreover, the amino-acid composition of the functional neighborhoods revealed a distinctive biochemical signature (**Fig. 5f**; Methods). Relative to random background, these neighborhoods were enriched for residues known to contribute to protein interaction interfaces, particularly aromatic residues (such as tryptophan, tyrosine) and arginine, which are characteristic of energetic “hot spots” clustered near the cores of binding interfaces.^11,35,36^ These analyses show that ReCLIP learns residue-centered neighborhoods that are structurally and functionally coherent, providing a biologically grounded and interpretable basis for its predictive performance.

### ReCLIP identifies clinically relevant interaction perturbations

To evaluate the clinical relevance of predicted interaction perturbations, we applied ReCLIP to clinically annotated variants from ClinVar and classified each variant by its predicted effect on protein interactions (disrupting, decreasing, increasing, or no effect; Methods). Across 144,837 missense variants evaluated over 500,083 interaction pairs, predicted interaction perturbations were significantly more frequent among pathogenic and likely pathogenic (PG/LP) variants than among benign and likely benign (BN/LB) variants (**Fig. 6a**). This enrichment was evident both at the level of individual variant-interaction pairs (135,951/201,021=67.6% versus 122,481/299,062=41.0%, Fisher’s exact test *P*<1e-200) and when effects were aggregated across variants (49,393/74,263=66.5% versus 52,490/116,487=45.1%, Fisher’s exact test *P*<1e-200). Applying more stringent class-specific probability thresholds further sharpened the separation (**Extended Data Fig. 2a**), supporting the use of ReCLIP predictions to prioritize missense variants with potential clinical relevance.

**Figure 6.**
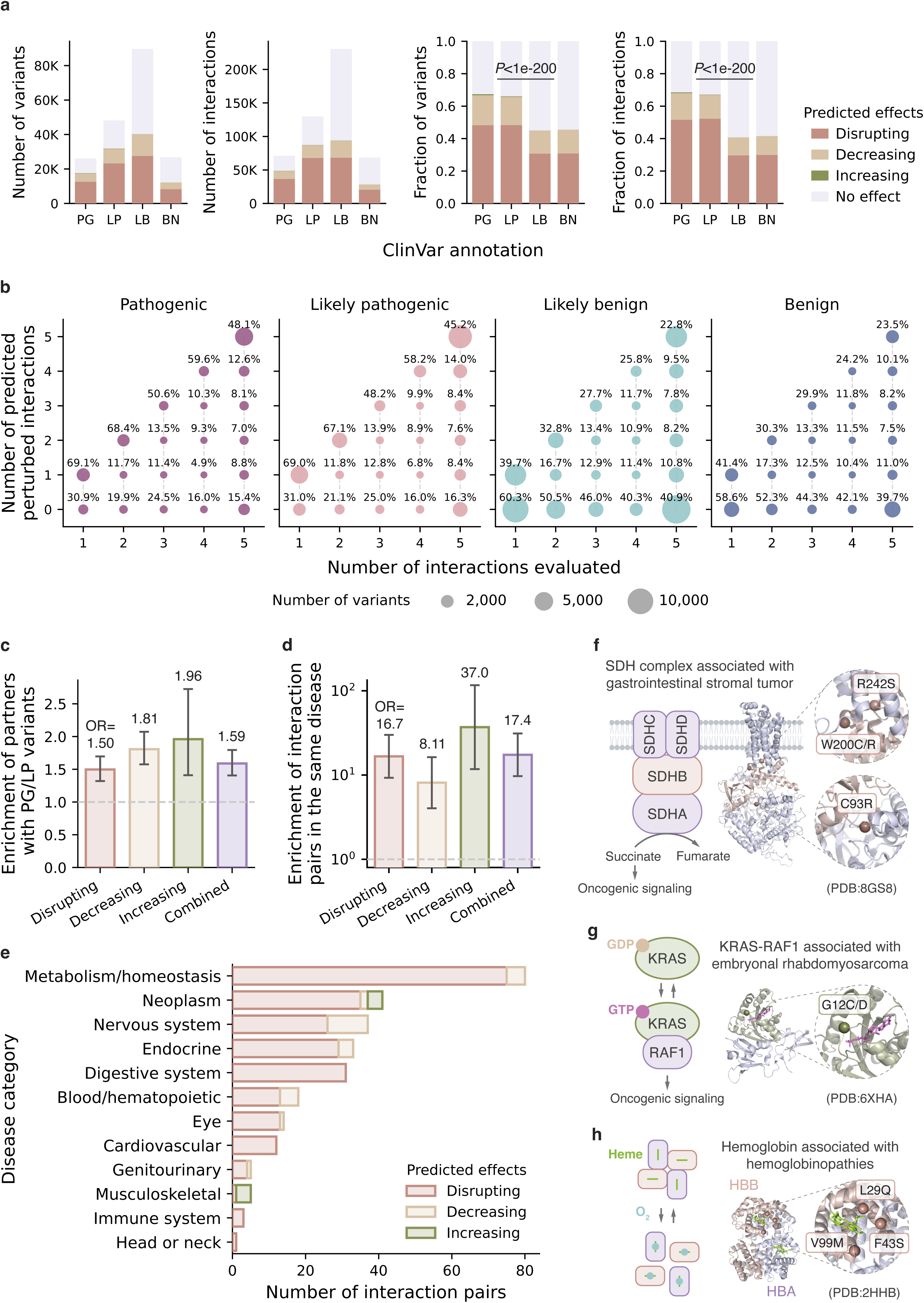
ReCLIP identifies clinically relevant interaction perturbations. **a**, Predicted interaction perturbations among clinically annotated ClinVar missense variants. Variants were evaluated across protein interactions and classified into four outcome classes (disrupting, decreasing, increasing, or no effect) based on ReCLIP predictions. Left, total numbers of variants and variant–interaction pairs, stratified by predicted interaction outcome and ClinVar annotation. Right, corresponding fractions of predicted perturbation classes across variants and interaction pairs. Predicted perturbations were enriched among pathogenic and likely pathogenic (PG/LP) variants relative to benign and likely benign (BN/LB) variants at both variant and interaction levels. Statistical significance was assessed using two-sided Fisher’s exact tests. **b**, Predicted perturbation frequencies across different numbers of interaction partners evaluated per variant. Bubble size indicates the number of variants in each category, and percentages indicate the fraction of variants predicted to perturb the corresponding number of interactions. PG/LP variants showed consistently higher perturbation frequencies than BN/LB variants across one to five interaction partners. **c**,**d**, Enrichment of perturbed interaction partners harboring PG/LP variants (**c**) and interaction pairs sharing disease phenotypes (**d**). Bars indicate odds ratios (ORs) for each perturbation class and for all perturbations combined relative to no-effect predictions; error bars represent 95% CIs. **e**, Distribution of perturbed interaction pairs sharing phenotype annotations across major disease categories. Disease categories were derived from first-level Human Phenotype Ontology (HPO) subclasses (HP:0000118). Bars are stratified by predicted perturbation class. **f-h,** Representative examples illustrating that ReCLIP links variant effects to plausible molecular mechanisms. **f,** Predicted interaction-disrupting variants in SDHB localize to interaction interfaces within the succinate dehydrogenase (SDH) complex, disruption of which is associated with succinate-driven oncogenic signaling. **g,** Predicted interaction-increasing KRAS G12 variants localize near the GTP-binding site, where stabilization of the GTP-bound state enhances RAF1 signaling. **h,** Predicted interaction-disrupting variants in hemoglobin localize near the heme pocket, where oxygen binding regulates subunit interactions and tetramer conformational transitions. Highlighted residues indicate PG/LP variants mapped onto experimentally resolved protein structures (PDB IDs: 8GS8, 6XHA, and 2HHB, respectively).

Because individual proteins often participate in multiple interactions, we examined perturbation patterns across up to five interaction partners per variant (Methods). PG/LP variants showed consistently higher perturbation frequencies than BN/LB variants across one to five partners (**Fig. 6b**). Among variants evaluated against five partners, 84.0% (23,565/28,058) of PG/LP variants were predicted to perturb at least one interaction, and nearly half (12,969/28,058) perturbed all five interactions (1.41- and 2.01-fold enrichments relative to BN/LB variants; **Fig. 6b**).

To control for protein- and interaction-specific context, we compared PG/LP and BN/LB variants within matched interaction pairs. Within the same interaction, PG/LP variants exhibited higher perturbation probabilities in the majority of cases (8,593 of 10,357 pairs, 83.0%), with a significantly higher overall perturbation probability than BN/LB variants (medians=0.68 versus 0.55, paired Wilcoxon signed-rank test *P*<1e-200; **Extended Data Fig. 2b**). These results indicate that the enrichment of predicted perturbations among PG/LP variants is not driven by protein- or interaction-level context, but reflects consistent residue-level differences in predicted molecular consequences.

We next assessed the biological relevance of perturbed interaction partners. Compared with predicted no-effect interactions, perturbed partners of PG/LP variants were enriched for proteins that also harbor PG/LP variants (OR = 1.59, Fisher’s exact test *P*=1.57e-14; **Fig. 6c**). Moreover, these interaction pairs were more likely to involve proteins with shared disease phenotypes (OR=17.4, Fisher’s exact test *P*=1.14e-46; **Fig. 6d**; Methods), suggesting that ReCLIP captures coordinated perturbations within functionally coherent, disease-associated interaction contexts. These associations spanned diverse disease categories, including metabolism and homeostasis, neoplasms, nervous system disorders, digestive disease, and hematopoietic phenotypes (**Fig. 6e**).

Representative examples illustrate how ReCLIP links variant effects to plausible molecular mechanisms. In mitochondrial metabolism, predicted interaction-disrupting variants in SDHB localized to interfaces within the succinate dehydrogenase (SDH) complex (**Fig. 6f**), whose disruption is known to impair mitochondrial respiration and promote metabolite-driven oncogenic signaling.^37^ In RAS signaling, ReCLIP predicted increased interaction between KRAS G12 variants and RAF1, despite the mutated residues being distal from the binding interface (**Fig. 6g**). This is consistent with allosteric mechanisms in which G12 substitutions increase the active GTP-bound state and enhance downstream RAF signaling.^38,39^ Similarly, in hemoglobin, predicted perturbations localized to residues near the heme pocket (**Fig. 6h**), reflecting the known allosteric coupling between oxygen binding and tetramer conformational transitions, through which variant effects can indirectly alter subunit interactions.^40,41^

Together, these analyses demonstrate that ReCLIP can distinguish clinically pathogenic from benign variants, identify disease-relevant perturbations across molecular interaction contexts, and provide mechanistic insight into how genetic variation alters protein interactions through both direct interface perturbation and non-local regulatory effects.

## Discussion

PLMs have advanced protein modeling by learning rich residue-level representations from sequence, yet most PLM-based approaches to PPIs rely on protein-level aggregation. As a result, local determinants of interaction specificity can become difficult to resolve, limiting the ability to distinguish context-dependent interaction outcomes. Here, we present ReCLIP, which explicitly models this context through learned functional neighborhoods and improves PPI prediction across diverse biological settings, spanning residue-level perturbations and distinct interaction contexts.

A central implication of our evaluation is that residue-centered context provides a useful organizing principle for modeling protein interactions. This design is particularly important for residue-level perturbation prediction, where the same protein pair can exhibit distinct outcomes depending on the specific residue being affected. Existing interaction-aware approaches partially capture this by presenting mutated protein pairs as input, yet ultimately compress the resulting representations into protein-level embeddings that may dilute residue-specific effects.^9^ Moreover, approaches that rely on comparing wild-type and mutant sequences are not applicable to perturbations that do not change primary sequence (such as PTMs). In contrast, ReCLIP learns functional neighborhoods centered on each residue, enabling direct modeling of interaction-relevant context and generalization to perturbations without requiring explicit sequence change. The pMHC results further highlight the ability of residue-centered modeling to generalize across diverse interaction contexts. Predicting peptide binding to MHC alleles requires extrapolating across extensive receptor polymorphism while preserving sensitivity to residue-level determinants of peptide recognition. The consistently high zero-shot performance of ReCLIP suggests that the model captures shared residue-level constraints relevant to pMHC binding rather than relying solely on allele-specific patterns. Collectively, these tasks span qualitatively distinct biological settings, establishing residue-centered interaction modeling as a general framework for protein binding that is not tied to a specific perturbation type or task formulation.

These findings complement emerging evidence that PLMs encode rich residue-level functional information and show that such information can be organized for protein interaction modeling. To evaluate the importance of residue-conditioned neighborhood selection, we constructed a control model in which functional neighborhoods were replaced with randomly sampled residues from the same protein, approximating protein-level aggregation. Predictive performance declined substantially (**Extended Data Fig. 3a**), indicating that the improvements achieved by ReCLIP depend on selective residue-centered context rather than global aggregation of sequence information.

The analysis of learned neighborhoods provides a biological rationale for these performance gains. Residues prioritized by ReCLIP are not uniformly distributed along the sequence, but form coherent neighborhoods that extend beyond immediate sequence adjacency and reflect broader structural and functional relationships. Importantly, these biologically coherent neighborhoods are also evident in clinically relevant settings. Predicted interaction perturbations were enriched among pathogenic variants and organized within interaction pairs sharing disease phenotypes, reinforcing that residue-centered interaction modeling captures coordinated perturbations within disease-associated molecular contexts rather than nonspecific interaction effects. This perspective provides a framework for understanding how genetic variation propagates through interaction networks to influence disease biology. More broadly, interaction-centered representations could complement existing variant interpretation frameworks by providing molecular context for variant impact and prioritizing therapeutically actionable perturbations, including variants directly targeted by small-molecule inhibitors such as KRAS G12C.^38,39^

Despite these advances, several limitations should be noted. First, the framework depends on pretrained PLM embeddings and therefore inherits biases and representational constraints from the underlying model. In this study, we employed embeddings from the final transformer layer of ESM-2, selected to align backbone capacity with the PLM-based baselines. Evaluation across alternative transformer layers yielded similar performance (**Extended Data Fig. 3b**), suggesting that ReCLIP is robust to layer selection, although systematic benchmarking across PLM architectures and training regimes remains an important direction for future work. Second, the curated datasets used for evaluation, although widely adopted, are inherently enriched for well-studied proteins and experimentally tractable perturbations. This bias is particularly evident for underrepresented outcome classes, such as interaction-increasing mutations and interaction-inhibiting PTMs, where predictive performance remains comparatively modest across all methods. Nevertheless, ReCLIP consistently achieves larger performance gains in these challenging settings, indicating robustness to the sparse and unevenly distributed observations that characterize many biological datasets. Finally, although the learned neighborhoods align with known structural and functional features, model attention weights should not be interpreted as direct evidence of causal importance. The prioritized residues provide informative hypotheses about interaction-relevant context but do not establish mechanistic causality. Determining which of these learned associations correspond to bona fide determinants of interaction specificity will require targeted functional validation.

In summary, our study supports a residue-centered view of protein interactions in which binding specificity and its perturbation are encoded in localized, structurally and functionally coherent neighborhoods. By explicitly modeling these neighborhoods and their partner-specific dependencies, ReCLIP provides a generalizable and interpretable framework for predicting protein interaction outcomes and linking residue-level perturbations to disease-relevant molecular contexts across diverse biological settings.

## Methods

### ReCLIP framework

ReCLIP is a residue-centered machine learning framework for modeling protein-protein interactions (PPIs) directly from sequence. ReCLIP takes as input a target protein sequence, an interaction partner sequence, and a queried residue position on the target protein. The queried residue is task-specific and may correspond to a mutation site, a PTM residue, or a peptide anchor position in a pMHC molecule.

ReCLIP integrates two complementary feature representations to model PPIs at residue resolution. The intra-protein representation encodes the queried residue together with its learned functional neighborhood in the target protein, capturing sequence-derived dependencies among residues. The inter-protein representation encodes the interaction partner conditioned on the queried residue, with partner residues weighted according to their relevance to the query. These representations are concatenated into a joint feature vector and used for supervised prediction of interaction outcomes.

### Intra-protein feature representation

ReCLIP employs self-attention patterns from a pretrained ESM-2 model to construct residue-centered representations of the target protein. ESM-2 is a transformer-based protein language model (PLM) whose self-attention weights encode context-dependent dependencies between residues, capturing biochemical, structural, and evolutionary relationships learned during large-scale pretraining. We adopted the 650M-parameter ESM-2 backbone (esm2_t33_650M_UR50D) to match the scale of the PLM-based baselines, ensuring that performance differences primarily reflected architectural differences rather than backbone scale. Residue embeddings (1,280 dimensions) and corresponding L×L self-attention matrices (up to 510×510 depending on input sequence length) were extracted from the final transformer layer (layer 33). Robustness to embedding layer selection was evaluated using alternative ESM-2 layers (**Extended Data Fig. 3b**).

For each queried residue, self-attention weights from the queried position to all other residues in the target sequence were extracted from all attention heads (n=20) of the final layer. Let *A*_!_(*i*, *j*) denote the attention weight assigned by attention head *h* from queried residue *i* to residue *j*. Within each head, residues were ranked by attention weight, and the top-*k* residues were selected to define the functional neighborhood:

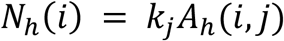

For each selected residue *j* ∈ *N*_!_(*i*), the corresponding hidden-state embedding from the same transformer layer was extracted. Let *z*(*j*) ∈ *R*^#^ denote the residue embedding for residue *j*, where *d*=1,280. Embeddings were concatenated in descending attention order to form a head-specific representation:

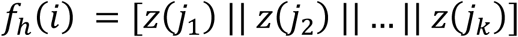

where *j*_1_, … , *j*_k_ denote residues ranked by attention weight. This rank-preserving concatenation retains the relative hierarchy of residue importance and preserves complementary neighborhood information captured by different attention heads, in contrast to pooling-based approaches that collapse this information into averaged representations.

The final intra-protein representation was constructed by concatenating head-specific neighborhood vectors across all attention heads:

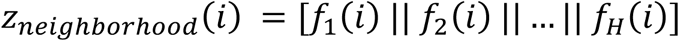

If fewer than *k* valid residues were available for a given head, zero-padding was applied to preserve output dimensionality. We selected *k* = 5 based on ablation experiments evaluating neighborhood sizes of *k* = 3, 5, 7, and 10, which showed comparable predictive performance but substantially increased computational cost for larger neighborhoods (**Extended Data Table 1**).

### Inter-protein feature representation

To model interaction-specific dependencies between the target and partner proteins, ReCLIP constructs a residue-conditioned representation of the interaction partner. Target and partner sequences were jointly encoded using MINT, a multimer-aware PLM built upon the ESM-2 architecture and trained to model interacting protein sequences.

For queried residues on the target protein, inter-protein (cross-chain) attention weights to partner residues were extracted from the MINT final layer (up to 1,020×1,020 depending on input sequence length). Let *A*_*h*_^*cross*^(*i*, *j*) denote the attention weight from queried residue *i* to partner residue *j* in attention head *h*. These weights were averaged across all 20 heads to reduce head-specific variability and normalized to define a consensus partner relevance distribution:

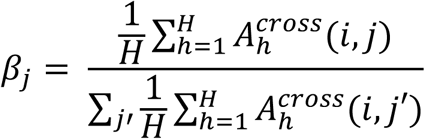

Let *z*(*j*) denote the hidden-state embedding of partner residue *j* from the same transformer layer. The residue-conditioned partner representation was computed as a weighted sum of partner residue embeddings:

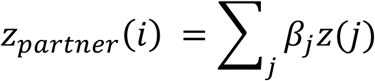

This formulation allows the same protein pair to generate distinct representations depending on the queried residue, enabling residue-specific modeling of interaction context.

### Joint interaction feature representation and supervised prediction

The final ReCLIP representation was constructed by concatenating intra-protein and partner-conditioned feature vectors:

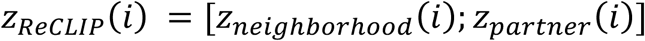

This joint feature representation was used as input to a gradient-boosted decision-tree classifier implemented using XGBoost. Multiclass tasks (e.g., mutation effect prediction) were trained using the multi:softprob objective, whereas binary tasks (e.g., PTM effect prediction and pMHC binding prediction) used binary:logistic.

XGBoost hyperparameters were fixed across tasks: n_estimators = 3500, learning_rate = 0.02, max_depth = 4, min_child_weight = 12, subsample = 0.85, colsample_bytree = 0.85, gamma = 1.0, reg_lambda = 10.0, and reg_alpha = 0.1. Hyperparameter optimization was performed exclusively on training partitions to prevent information leakage.

### Mutation-induced PPI perturbation

*Dataset.* Mutation effect prediction data were curated from the IntAct database.^25^ We retained single amino acid substitutions with annotated effects on protein binding and mapped annotations into four outcome classes: disrupting, decreasing, increasing, and no effect. Entries were deduplicated by mutation, target protein, interaction partner, and outcome label, and mutation-interaction pairs with conflicting annotations were excluded. Records involving multiple interaction partners were expanded into pairwise target–partner interactions, followed by secondary deduplication and conflict resolution. Target and partner sequences were mapped to human UniProt/Swiss-Prot reference sequences,^42^ and entries lacking valid sequence mappings were removed. Mutant sequences were generated by applying the annotated amino acid substitution at the corresponding position. The final dataset comprised 56,429 mutation–interaction pairs, including 13,780 disrupting, 9,069 decreasing, 2,228 increasing, and 31,352 no-effect interactions.

*Prediction*. Prediction of mutation-induced PPI perturbation was formulated as a four-class classification task. For each instance, ReCLIP used the mutated target sequence, mutation position, and partner sequence to generate the joint interaction representation. Model performance was evaluated using 10-fold stratified cross-validation while preserving class proportions across folds. Evaluation metrics included multiclass area under the curve (AUROC) and area under the precision-recall curve (AUPRC), accuracy, balanced accuracy, macro-precision, macro-recall, and macro-F1 score. To evaluate ranking performance, predictions were ranked separately for each class according to the class-specific probability produced by the classifier, treating each class in a one-vs-rest manner. Precision was computed at multiple Top-K thresholds for each class, and the final ranking metric was reported as macro-precision averaged across the four classes.

### PTM-regulated PPI perturbation

*Dataset* PTM effect prediction data were curated from the PTMint database.^30^ We retained phosphorylation events experimentally annotated as enhancing or inhibiting binding. Records were deduplicated by modification site, target protein, interaction partner, and outcome label. The final dataset comprised 3,052 PTM-interaction pairs, including 2,084 enhancing and 968 inhibiting instances.

*Prediction* Prediction of PTM-regulated PPI perturbation was formulated as a binary classification task (enhancing versus inhibiting). Each instance was defined by the target protein sequence, modification site, and interaction partner sequence, and interaction representations were generated using the modification site as the queried residue. Because PTMs do not alter the primary amino acid sequence, methods requiring explicit comparison between wild-type and mutant sequences (MINT, AlphaMissense, and PrimateAI-3D) were not applied to this task. Performance was evaluated using 10-fold stratified cross-validation. Evaluation metrics were consistent with those used for mutation-induced perturbation prediction, including AUROC, AUPRC, balanced accuracy, precision, recall, and F1 score.

### pMHC binding

*Dataset.* pMHC interaction datasets were obtained from previously curated resources assembled by SWING.^17^ The original datasets were derived from the NetMHCpan v4.1^32^ and NetMHCIIpan v4.2^43^ benchmark collections and comprised experimentally supported pMHC interactions spanning multiple HLA alleles. MHC sequences were obtained from the MHC Restriction Ontology,^44^ and alleles were grouped according to functional similarity using MHCCluster v2.0.^45^ The class I training set included eight HLA alleles: HLA-B*15:03, HLA-C*07:02, HLA-A*25:01, HLA-C*08:02, HLA-B*07:02, HLA-A*02:11, HLA-A*11:01, and HLA-B*40:01. Held-out alleles were divided into functionally proximal (HLA-A*02:02, HLA-B*40:02, and HLA-C*05:01) and functionally distal (HLA-A*32:01, HLA-C*03:03, and HLA-B*38:01) sets. For mixed-class analyses, the class I training data were combined with curated class II datasets comprising DRB1*01:01, DRB1*03:01, DRB1*04:01, DRB1*07:01, and DRB1*15:01.

*Prediction* pMHC binding prediction was formulated as a binary classification task (binding versus not binding). Interaction representations were generated for each pMHC pair, with the peptide sequence used as the target sequence, its central position as the queried residue, and the MHC binding groove as the interaction partner. Evaluation was performed under strict zero-shot settings in which test alleles were excluded from training. The class I model was evaluated on proximal and distal held-out alleles to assess generalization across varying degrees of MHC similarity, and the mixed-class model was evaluated on the same class I held-out alleles to assess robustness to heterogeneous training data. Evaluation metrics were consistent with those used for mutation and PTM tasks.

### Benchmark models

ReCLIP was benchmarked against state-of-the-art PLM-based protein interaction models and single-protein variant effect predictors. MINT^9^ is an interaction-aware PLM that jointly encodes interacting protein sequences and generates pair-conditioned representations through cross-chain attention. SWING^17^ is a sequence-window-based interaction model that represents PPIs using local sequence neighborhoods centered on fixed sequence windows. A BERT-based PLM was included as a global sequence-embedding baseline, in which proteins were independently encoded and pooled into fixed-length representations. For mutation-induced perturbation prediction, AlphaMissense^26^ and PrimateAI-3D^27^ were included as single-protein-level variant effect predictors that do not explicitly model interaction partners. For binary mutation effect prediction, we additionally compared ReCLIP with recent PLM-based interaction models developed for binary classification, including PLM-Interact^29^ and eSIG-Net,^28^ evaluated on the disrupting-versus-no-effect subset of the IntAct dataset. All baseline models were implemented using the recommended settings or publicly available configurations described in the original publications.

### Bootstrap significance testing

Statistical significance of performance differences between models was assessed using paired bootstrap resampling. For each evaluation metric, predictions from ReCLIP and the strongest competing baseline were compared on identical test instances using 200 paired bootstrap resamples of the test set generated by sampling with replacement. For bootstrap iteration *b*, the metric difference was defined as:

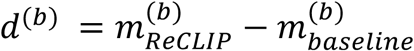

The bootstrap distribution of *d*(b) was summarized by its mean and 95% confidence interval (CI: 2.5^th^-97.5^th^ percentiles). Statistical significance was assessed by approximating the bootstrap difference distribution as Gaussian and computing the standardized test statistic:

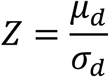

where *μ*_d_and *σ*_d_denote the mean and standard deviation of the bootstrap differences, respectively. Two-sided *P*-values were computed from the standard normal distribution.

### Analysis of residue-centered functional neighborhoods

To characterize the biological properties of residue-centered representations learned by ReCLIP, we analyzed the sequence, structural, and functional organization of attention-derived residue neighborhoods in the mutation perturbation task. ReCLIP-prioritized residue neighborhoods were defined as the top five residues receiving the highest self-attention weights from the queried residue within each of the 20 attention heads. Matched random residue sets were generated independently for each queried residue by sampling an equal number of residues from the corresponding protein sequence. Unless otherwise specified, all analyses were performed in a paired manner between prioritized and random residue sets associated with the same queried residue.

#### Sequence distance

Sequence distance was defined as the absolute difference in residue index between the queried residue and each prioritized or random residue along the primary protein sequence. For each queried residue, the mean sequence distance was computed across prioritized residues aggregated over 20 attention heads and compared with matched random residue sets. In total, 19,547 queried residues across 3,202 proteins were included in the analysis (n = 19,547 paired observations). Differences were assessed using two-sided paired Wilcoxon signed-rank tests.

#### Structural proximity

Structural proximity was quantified using residue-to-residue distances derived from AlphaFold-predicted structures.^46^ Distances were computed as the mean Euclidean distance between the Cα atom of the queried residue and those of prioritized or random residues. Analyses were restricted to high-confidence structures, defined as proteins with predicted local distance difference test (pLDDT) ≥ 70 for more than 80% of residues in the protein. Queried residues with at least one prioritized residue mapped to the corresponding structure were retained, resulting in 8,466 queried residues across 965 proteins (n = 8,466 paired observations). To assess non-local structural organization independently of sequence adjacency, analyses were additionally repeated after restricting residue neighborhoods to those separated by more than 10 amino acids in primary sequence. Differences were assessed using two-sided paired Wilcoxon signed-rank tests.

#### Functional annotation enrichment

Functional enrichment was evaluated using UniProtKB functional site annotations.^42^ Prioritized residues were intersected with annotated functional sites, including catalytic sites, binding sites, post-translationally modified residues, and disulfide-bonded cysteines. Enrichment was quantified by comparing annotation frequencies between prioritized and background residues using two-sided Fisher’s exact tests. In total, 56,360 of 1,269,100 (4.4%) prioritized residues overlapped annotated functional sites, compared with 125,650 of 5,465,707 (2.3%) background residues across 2,369 proteins. Analyses were additionally performed separately for catalytic sites, binding sites, modified residues, and disulfide-bonded cysteines.

We further evaluated the structural proximity of prioritized residues to annotated functional sites using AlphaFold-predicted structures. For each queried residue, Euclidean distances to the nearest annotated site were averaged across associated prioritized (or random) residues. Analyses were performed both across all annotations (n = 4,059 paired observations) and separately for catalytic sites (n = 953), binding sites (n = 1,998), modified residues (n = 3,441), and disulfide-bonded cysteines (n = 719). Consistent structural quality-control filters were applied as described in the structural proximity analysis. Differences were assessed using two-sided paired Wilcoxon signed-rank tests.

#### Domain coherence

Domain-level organization was evaluated using InterPro annotations.^47^ For each queried residue, domain coherence was defined as whether all five residues prioritized by an attention head co-localized within the same annotated domain interval as the queried residue. Analyses were restricted to proteins containing at least one annotated domain interval and queried residues located within annotated domains. In total, 190,209 of 229,340 (82.9%) prioritized residue neighborhoods exhibited domain coherence, compared with 54,108 of 229,340 (23.6%) random residue sets across 1,872 proteins. Statistical significance was assessed using two-sided Fisher’s exact tests.

#### Amino-acid composition

Amino-acid composition was analyzed across the 20 canonical amino acids. Residue counts were aggregated across prioritized residues from all 20 attention heads and compared with matched random residues sampled from the corresponding protein sequences (19,547 queried residues across 3,202 proteins). Amino acids were ranked according to their relative enrichment in prioritized versus random residues.

#### Analysis of ClinVar variants

ClinVar variants were obtained from the UCSC Table Browser^48^ ClinVar SNVs track. Analyses were restricted to single amino acid substitutions with clinical significance corresponding to pathogenic (PG), likely pathogenic (LP), benign (BN), or likely benign (LB) annotations. Variants were mapped to UniProt protein sequences and retained only if the annotated amino acid substitution matched the corresponding UniProt residue and position. Protein interaction partners were retrieved from the HINT database,^49^ which compiles high-quality experimentally supported physical interactions. For each protein harboring ClinVar variants, variant effects were evaluated against up to five interaction partners.

Interaction perturbations were inferred using ReCLIP trained on the IntAct dataset under a four-class classification setting (disrupting, decreasing, increasing, and no effect). For each ClinVar variant-interaction pair, the mutant protein sequence, variant position, and interaction partner sequence were used to generate residue-centered interaction representations and to predict interaction outcomes. A total of 591 variant-interaction pairs overlapping between the ClinVar and IntAct datasets were excluded from downstream analyses. The final dataset comprised 144,837 ClinVar missense variants (19,045 PG, 35,347 LP, 20,811 BN, and 69,634 LB variants) evaluated across 500,083 interaction pairs.

For the disease phenotype analysis in Fig. 6d, phenotype annotations were obtained from Human Phenotype Ontology (HPO) terms associated with ClinVar entries in the original UCSC ClinVar annotation table. Interaction pairs were considered to share a disease phenotype if the two interacting proteins were annotated with at least one common HPO term. For the disease category analysis in Fig. 6e, HPO annotations were mapped to first-level subclasses under HP:0000118 (“Phenotypic abnormality”) to define broad disease categories.

#### Data availability

The mutation-induced PPI perturbation dataset was obtained from the IntAct database^25^ https://www.ebi.ac.uk/intact/download/datasets#mutations. The PTM-regulated interaction dataset was downloaded from PTMint^30^ https://ptmint.sjtu.edu.cn/. pMHC datasets were derived from previously curated datasets assembled by SWING^17^ https://github.com/jishnu-lab/SWING. Protein sequences and functional annotations were obtained from UniProtKB/Swiss-Prot^42^ https://www.uniprot.org/. Predicted and experimentally resolved protein structures were obtained from the AlphaFold Protein Structure Database^46^ https://alphafold.ebi.ac.uk/ and the RCSB Protein Data Bank (PDB)^50^ https://www.rcsb.org/, respectively. Domain annotations were obtained from InterPro^47^ https://www.ebi.ac.uk/interpro/. ClinVar variants and annotations were obtained from the UCSC Table Browser^48^ ClinVar SNVs (clinvarMain) table under the ClinVar Variant track https://genome.ucsc.edu/cgi-bin/hgTables. Protein interactions used for ClinVar interaction perturbation inference were retrieved from the HINT^49^ Homo sapiens binary high-quality interaction dataset https://hint.yulab.org/.

## Code availability

All code used for data processing, model training, benchmarking, and statistical analyses are publicly available at https://github.com/SiweiLab/ReCLIP.

---

**Extended Data Figure 1.**
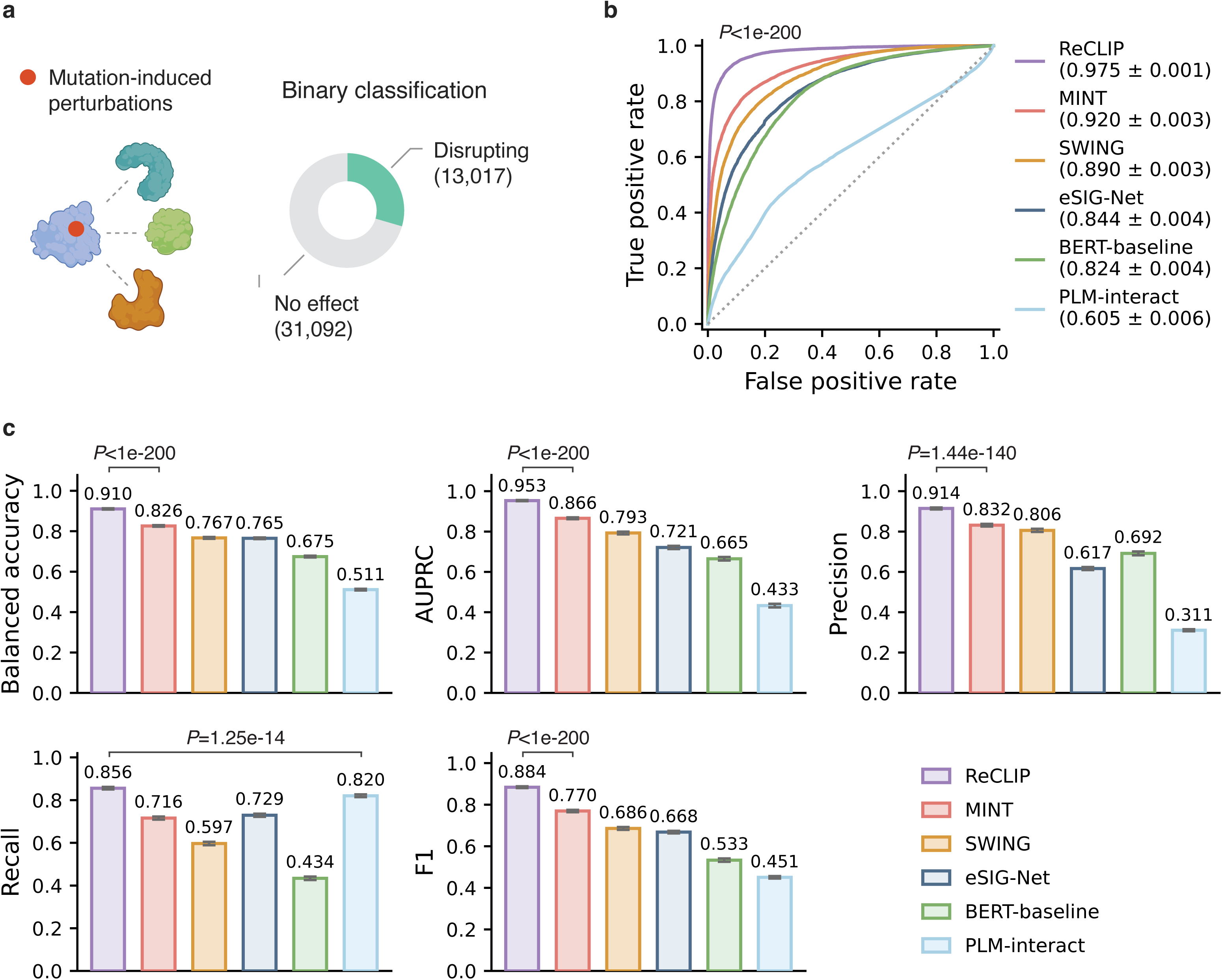
ReCLIP maintains robust performance across different task formulations. **a**, Overview of the mutation-induced interaction perturbation prediction reformulated as a binary classification problem (disrupting versus no effect). **b,c**, Performance comparison of ReCLIP with baseline methods, including two additional PLM-based interaction models designed for binary prediction. ReCLIP achieves the highest overall AUROC (**b**) and consistently outperforms comparison methods across evaluation metrics (**c**). Evaluation metrics are reported as mean with 95% CIs computed from 200 bootstrap resamples; statistical significance was assessed from the bootstrap distribution of metric differences between ReCLIP and the second-best method.

**Extended Data Figure 2.**
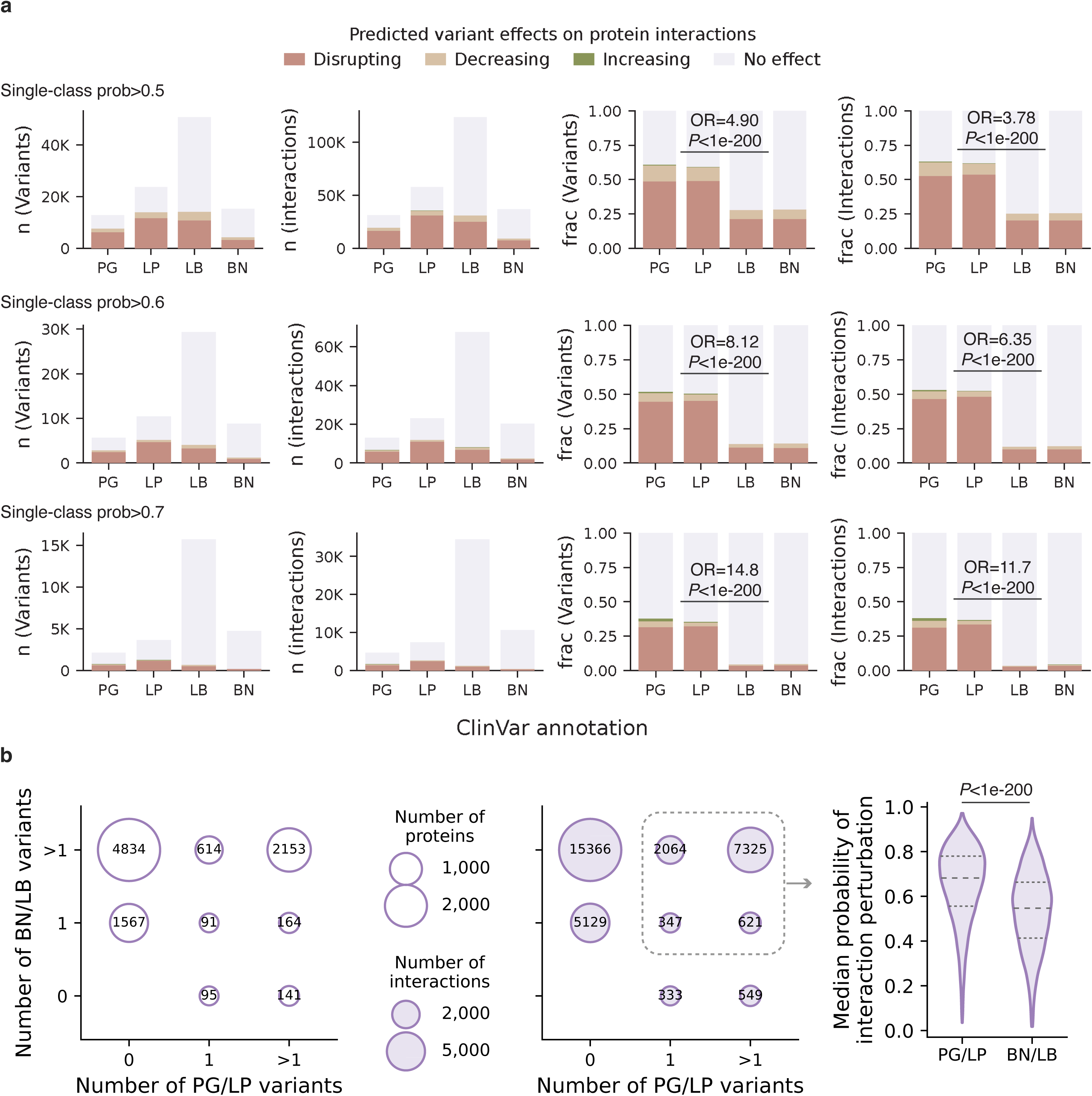
ReCLIP prioritizes pathogenic variants. **a**, Predicted interaction perturbations for ClinVar missense variants under different single-class probability thresholds. Variants were classified into four outcome classes (disrupting, decreasing, increasing, or no effect) based on ReCLIP predictions. Rows correspond to increasingly stringent thresholds requiring the predicted probability of a single interaction outcome class to exceed 0.5, 0.6, or 0.7. Left two columns, total numbers of variants and variant-interaction pairs, stratified by predicted interaction outcome and ClinVar annotation. Right two columns, corresponding fractions of predicted perturbation classes across variants and interaction pairs. Increasing probability thresholds sharpened the separation between PG/LP and BN/LB variants. Odds ratios (ORs) and Fisher’s exact test P values are shown.**b**, Comparison of PG/LP and BN/LB variants within matched interaction contexts. Left, distribution of proteins stratified by the numbers of PG/LP and BN/LB variants. Middle, corresponding distribution of interaction pairs. Right, comparison of median predicted perturbation probabilities for PG/LP and BN/LB variants within the same interaction pairs. Statistical significance was assessed using a two-sided Wilcoxon signed-rank test.

**Extended Data Figure 3.**
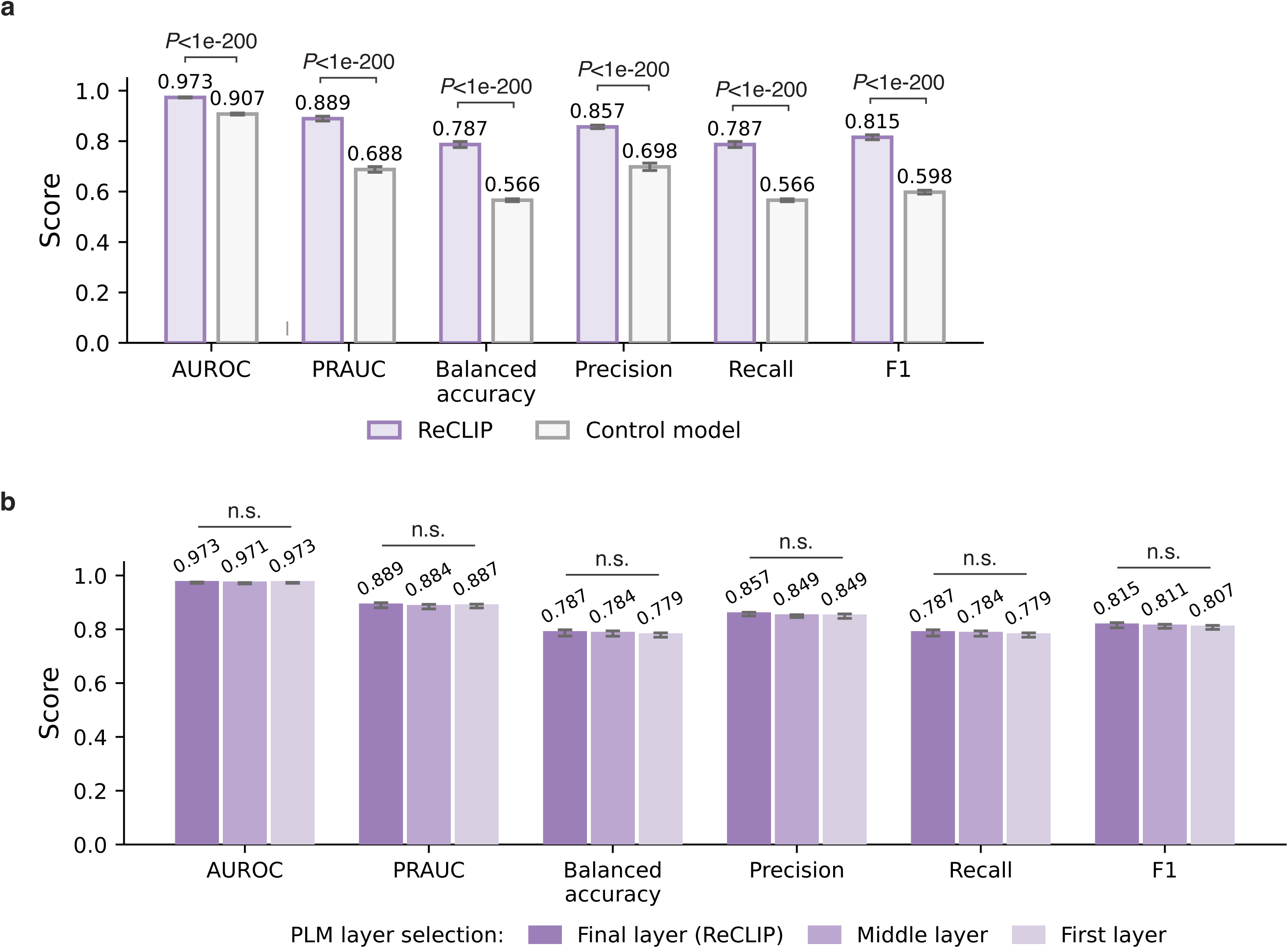
ReCLIP captures functional residue context and is robust to PLM layer selection. **a,** Comparison of ReCLIP with a control model in which selected functional neighborhoods are replaced with randomly sampled residues. ReCLIP outperforms the control model across evaluation metrics, supporting the importance of residue-conditioned neighborhood selection. **b,** Performance of ReCLIP using embeddings from different ESM-2 transformer layers, including the final layer (layer 33), the middle layer (layer 24), and the first layer. ReCLIP maintains consistent performance across layers, indicating robustness to embedding layer choice. For both panels, evaluation metrics are reported as the mean; error bars indicate 95% CIs computed from 200 bootstrap resamples. Statistical significance was assessed from the bootstrap distribution of metric differences between ReCLIP and the comparison model. n.s., not significant (*P* > 0.05).

**Extended Data Table 1.** Effect of neighborhood size on ReCLIP performance and computational cost. ReCLIP was evaluated using neighborhood sizes of *k* = 3, 5, 7, and 10 on the mutation-effect and PTM-effect prediction tasks. Predictive performance is reported as AUROC (mean ± 95% confidence interval [CI]), with CIs estimated from 200 bootstrap resamples. Runtime corresponds to the average computational time per cross-validation fold. Larger neighborhoods yielded only modest performance gains while substantially increasing computational cost.

|  | Mutation effect prediction |  | PTM effect prediction |  |
| --- | --- | --- | --- | --- |
| Neighborhood size (*k*) | AUROC ± 95% CI | Runtime per fold (s) | AUROC ± 95% CI | Runtime per fold (s) |
| 3 | 0.972 ± 0.001 | 211.8 | 0.820 ± 0.014 | 15.6 |
| 5 | 0.973 ± 0.001 | 316.5 | 0.823 ± 0.015 | 21.5 |
| 7 | 0.974 ± 0.001 | 398.3 | 0.824 ± 0.015 | 27.7 |
| 10 | 0.975 ± 0.001 | 546.5 | 0.825 ± 0.013 | 37.2 |

